# The receptor binding domain of SARS-CoV-2 spike is the key target of neutralizing antibody in human polyclonal sera

**DOI:** 10.1101/2020.08.21.261727

**Authors:** Tara L. Steffen, E. Taylor Stone, Mariah Hassert, Elizabeth Geerling, Brian T. Grimberg, Ana M. Espino, Petraleigh Pantoja, Consuelo Climent, Daniel F. Hoft, Sarah L. George, Carlos A. Sariol, Amelia K. Pinto, James D. Brien

## Abstract

Natural infection of SARS-CoV-2 in humans leads to the development of a strong neutralizing antibody response, however the immunodominant targets of the polyclonal neutralizing antibody response are still unknown. Here, we functionally define the role SARS-CoV-2 spike plays as a target of the human neutralizing antibody response. In this study, we identify the spike protein subunits that contain antigenic determinants and examine the neutralization capacity of polyclonal sera from a cohort of patients that tested qRT-PCR-positive for SARS-CoV-2. Using an ELISA format, we assessed binding of human sera to spike subunit 1 (S1), spike subunit 2 (S2) and the receptor binding domain (RBD) of spike. To functionally identify the key target of neutralizing antibody, we depleted sera of subunit-specific antibodies to determine the contribution of these individual subunits to the antigen-specific neutralizing antibody response. *We show that epitopes within RBD are the target of a majority of the neutralizing antibodies in the human polyclonal antibody response*. These data provide critical information for vaccine development and development of sensitive and specific serological testing.

## Main

<u>S</u>evere <u>a</u>cute <u>r</u>espiratory <u>s</u>yndrome <u>co</u>rona<u>v</u>irus 2 (SARS-CoV-2) was initially identified in patients with severe pneumonia in Wuhan, China in December of 2019. Due to its initial zoonotic transmission and human to human spread within an immunologically naïve population, it has since caused over 4 million confirmed cases and over 790,000 deaths worldwide (WHO 2020), with approximately 30% of all cases occurring in the United States as of July 15^th 1–5^. Infection with SARS-CoV-2 can result in a range of states from asymptomatic to symptomatic, with symptomatic cases ranging from mild non-specific symptoms, like malaise, to severe pneumonia and multiple organ failure ^1–3,5,6^.

SARS-CoV-2 is a positive sense, single stranded, enveloped RNA virus with a ~29 kb genome that is virologically similar to the enzoonotic beta-coronaviruses SARS-CoV and MERS-CoV. The SARS-CoV-2 genome encodes 16 non-structural proteins and 4 structural proteins: spike (S), nucleocapsid (N), envelope (E), and membrane (M). The coronavirus N protein functions by interacting with viral RNA to form the ribonucleoprotein, while E and M function in virion assembly and budding ^7–10^. Spike is a homotrimeric transmembrane protein that is comprised of two subunits per monomer, S1 and S2 that are responsible for binding the host cell receptor and viral fusion, respectively. Similarly to the human coronavirus NL-63 and SARS-CoV, SARS-CoV-2 spike uses human angiotensin converting enzyme 2 (ACE2) to gain entry into target cells ^8–10^. Specifically, the S1 subunit of SARS-CoV-2 contains the receptor binding motif (RBM) within the receptor binding domain (RBD) that makes direct contact with the ACE2 receptor for receptor-mediated entry ^9–11^. Important to note for antibody structural determinants, the pre-fusion confirmation of the trimeric spike has a range of states that are described as “up” or “down” based on the angle of RBD within S1. For a virion to be able to interact with ACE2 and gain entry into host cells, RBD must be in the “up” conformation between 50° and 70° that represents a receptor-binding active state ^10,12^. When the interaction between RBD and ACE2 is disrupted, the entry of SARS-CoV-2 into susceptible cells is blocked ^13,14^.

Spike is known to be a major antibody antigenic determinant for both MERS-CoV and SARS-CoV ^15^ that leads to the generation of protective immune responses including the production of highly neutralizing antibodies ^16–18^. Targets for these antibodies within spike include both conformation dependent and linear epitopes of RBD and the S2 fusion peptide. These neutralizing antibodies are proposed to block RBD-ACE2 receptor interactions or prevent S2 fusion with host membranes ^19,20^. Spike being a major antigenic determinant for the antibody response against closely related beta-coronaviruses contributes to our hypothesis that the neutralizing polyclonal antibody response to SARS-CoV-2 will target spike and its sub-domains.

## Results

### Antigenic variation of SARS-CoV-2 human isolates

To determine the current antigenic variation and display that variation within the structure of SARS-CoV-2 spike we interrogated 34,756 SARS-CoV-2 genomes derived from human samples available from GISAID on June 15, 2020. The spike homotrimer contains multiple subunits, including S1 and S2, both of which contain a total of 22 glycosylated residues which can affect spike protein folding, receptor interactions and potentially block antibody recognition and are represented as lollipops in the schematic (Figure 1A). The S1 subunit (residues 13-303) of spike contains the N terminal domain (NTD), C terminal domain (CTD), the receptor binding domain (RBD, residues 319-541), and the receptor binding motif (RBM, residues 437-508). The S2 subunit contains the fusion peptide (FP, residues 788-806), and heptad repeat 1 and 2 (HR1, residues 920-970, HR2, residues 1163-1202), the transmembrane domain and cytoplasmic tail. In our analysis of naturally occurring amino acid (aa) variation, low quality sequences determined by gaps or ambiguous nucleotides >50nt were removed (−497 sequences). The 34,259 remaining sequences were translated and aligned using MUSCLE (Supplemental file 1), then duplicate sequences were removed. This resulted in a multiple sequence alignment of 1273 amino acids (aa), with 4,065 unique aa sequences and an overall pairwise identity of 97.5%. The prevalence of aa variation per site and aa conservation was determined using the sequence variation tool (viprbrc) ^21^ and the CONSURF server ^22^, respectively. The level of aa variation was measured by calculating the aa frequency at each position within the multiple sequence alignment, then Shannon entropy was used to define aa conservation using data on the 20 potential aa present. These conservation scores were broken down into 6 discrete color coded categories with a score of 6 being most variable, representing >1,000 mutations at that site, and 1 being most highly conserved with 0-1 mutations per site. The aa conservation was then displayed in the context of the spike pre-fusion trimer (PDB:6VSB)^23^ to represent exposure to the human antibody response, where chain A displays the RBD within the up position and the B and C chains display the RBD in the down position (Figure 1B). The spike trimer color coded for aa variation is located next to the spike protein where the subunits are color coded with S1 NTD in cyan, RBD in dark green, and S2 in light green. From the aa sequence variation analyses, we observed the well documented G614D variation, which may have a fitness advantage. We also observed an additional 422 positions that contained a range of variation from 51/4065 (1.25%) to 2/4065 (0.05%). Once the aa variation was mapped onto the trimer structure, we observed that the greatest level of aa variation is found within the S1 NTD (94.2% identity), while the lowest level of aa variation is within the RBD (96.7% identity) and S2 (99.6%). The low level of aa variation within RBD was also recently described by Starr et al ^24^. Our data, in addition to that of Starr et al, indicate that overall the RBD and S2 domains are highly conserved and are currently genetically stable targets for vaccine and therapeutic intervention.

**Figure. 1.**
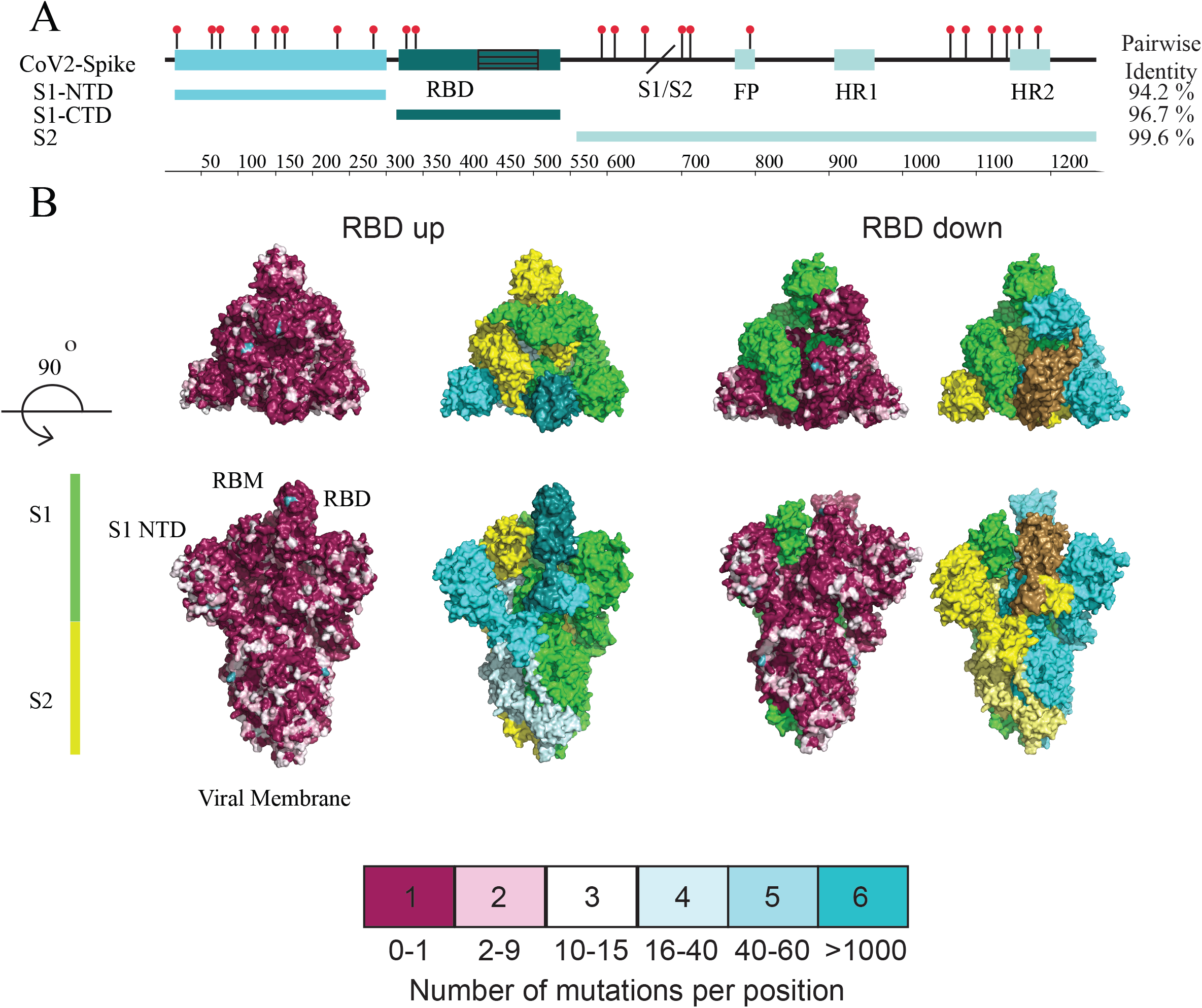
Antigenic variation of SARS-CoV-2 human isolates. (A)Schematic of the full-length SARS-CoV-2 spike protein with the S1 and S2 highlighted. S1 is divided into the N-terminal domain, and the C-terminal domain which contains the receptor binding domain (RBD) subunit in dark green with the receptor binding motif displayed using black hashed lines. The separation between S1and S2 is represented by a slash line. S2 contains the fusion peptide (FP) and heptad repeat one and two (HR1 and HR2). The spike protein is glycosylated at 22 residues with the location of each glycosylation indicated by a lollipop in the primary structure. (B) To determine the relative level of amino acid variation (diversity) within the spike protein of currently circulating SARS-CoV-2 strains, all high-quality sequences between Dec 2019 and June 2020 derived from humans were downloaded from GISAID, translated and aligned. Using the 4065 unique SARS-CoV-2 human derived spike sequences we calculated the positional percent diversity for all 1288 aa positions. The multiple sequence alignment identified 568 of 1288 positions which had at least one aa variant, where the number of variants ranged from 1-1277 for position 614. This sequence variation was displayed on the cryoEM reconstruction of the spike trimer (PDB:6VSB) where the surface reconstruction is colored according to 6 discrete groups, with a score of 1 being highly conserved (0-1 mutation per position) to being highly diverse with a score of 6 (>1000 mutations per position). The color coded bar describes the corresponding color for each range of mutations. Next to each aa variation coded structure is the cryoEM trimer structure with the individual trimers color coded to allow orientation. The forward facing trimer for the RBD up is color coded by subdomain, with RBD up being dark cyan, S1 as cyan, and S2 as pale green. The RBD down trimer is color coded with RBD down as brown, S1 as gold and S2 as pale yellow. We observed naturally occurring aa variations are less within the RBD as noted by the high level of purple colored Ca residues, and greatest aa variations within the S1-NTD as indicated by the white and green color residues.

We next wanted to investigate the specificity and immunodominance of the polyclonal antibody response to the subunits of spike due to its potential as a target for the development of vaccines and therapeutics. This concept is based upon vaccines developed against SARS-CoV and MERS-CoV ^15,25,26^ and the strongly neutralizing monoclonal antibodies that have been identified as spike specific ^27–32^. To address specificity and immunodominance, we analyzed serum samples from 16 laboratory confirmed SARS-CoV-2 infection cases as determined by qRT-PCR, with 11 of the 16 subjects being admitted to the hospital. The SARS-CoV-2 positive serum samples analyzed were obtained from patients in Saint Louis, MO, Cleveland, OH, and San Juan, PR, with no subjects succumbing to infection. The median age of the individuals is 59.1 years, with 7 males and 9 females. Specific demographics regarding the subjects are listed in Table 1. The cohort controls were collected prior to the emergence of SARS-CoV-2 (2015-2018) from previous studies conducted at Saint Louis, MO, Cleveland, OH, and San Juan, PR, and had a similar age and sex distribution.

**Table 1.** Demographics of SARS-CoV-2^+^ patient cohort. Demographics of the 16 patient cohort include sample number, sex, age, and day post laboratory confirmed SARS-CoV-2 positive test.

| **Samples Number** | **Sex** | **Age** | **Day Post Test** |
| --- | --- | --- | --- |
| 1 | F | 72 | 27 |
| 2 | F | 79 | 22 |
| 3 | M | 79 | 23 |
| 4 | M | 57 | 24 |
| 5 | F | 66 | 15 |
| 6 | F | 65 | 21 |
| 7 | F | 79 | 15 |
| 8 | M | 77 | 30 |
| 9 | M | 49 | 26 |
| 10 | M | 70 | 16 |
| 11 | F | 46 | 30 |
| 12 | F | 29 | 30 |
| 13 | M | 60 | 30 |
| 14 | F | 21 | 30 |
| 15 | M | 44 | 30 |
| 16 | F | 52 | 30 |

### Characterization of polyclonal IgG binding to SARS-CoV-2 Spike subunits

To investigate and quantify the IgG response to SARS-CoV-2, we performed ELISA assays using serum from 16 SARS-CoV-2^+^ subjects and 28 SARS-CoV-2 negative control subjects. We serially diluted sera from 1:50 to 1:64,000 as four-fold dilutions and evaluated binding to recombinant S1, S2, and RBD (Figure 2A-C). Polyclonal sera from all 16 SARS-CoV-2 subjects showed IgG binding to each spike subunit by ELISA. IgG reactivity to the S1 subunit, which contains the RBD, ranged in optical density (OD) from 1.4 to 3.4, while the 28 control subjects had an OD range from 0.13 to 2.2 at the highest concentration tested. Four subjects were responsible for the majority of the ELISA binding to S1 within the control group (Figure 2A). The overlap of these four negative subject samples with the SARS-CoV-2^+^ subjects suggests that there is antibody cross-reactivity to the SARS-CoV-2 S1 protein, most likely due to prior human coronavirus (HCoV) infections (NL63, HKU1, OC43, 229E). However, the focus of our study is to functionally define the key targets of the neutralizing antibody response to SARS-CoV-2, and further studies would need to be completed to define the nature of the cross-reactive response.

**Figure. 2.**
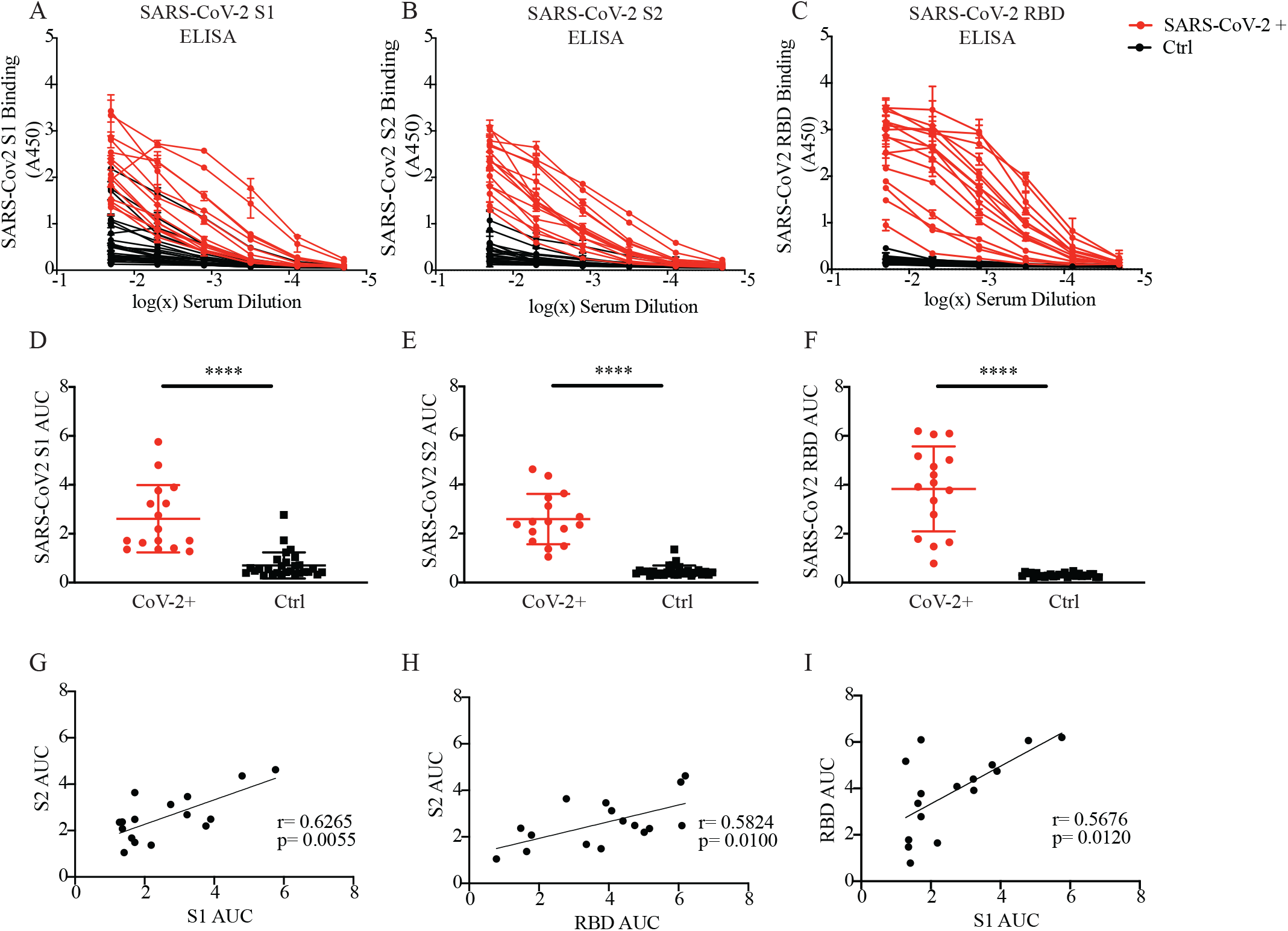
Characterization of polyclonal IgG binding to SARS-CoV-2 Spike subunits. Binding of polyclonal IgG from SARS-CoV-2 qRT-PCR positive patients and negative controls (ctrl) to recombinant SARS-CoV-2 proteins was determined by indirect ELISA. Binding curves derived from optical density at A450 for each subject are shown for (A) S1, (B) S2, and (C) RBD. Red lines indicate binding from SARS-CoV-2 + subjects and black lines indicate binding curves from ctrls. The total area under the curve (AUC) per subject was quantified from the corresponding binding curve and is shown for (D) S1 (E) S2, and (F) RBD. Similarly, red is indicative of SARS-CoV-2+ subjects and black represents the ctrls. Data are representative of n=16 for SARS-CoV-2 + and n=28 for ctrls. Statistical significance was determined by student’s 2-tailed t test *(*p* = 0.03), **(*p* = 0.002), ***(*p* = 0.0002), and ****(*p* < 0.0001). Immunodominance of the S subunits, in the context of the polyclonal antibody response, was probed by one-tailed spearman correlation analysis of the AUC values from the SARS-CoV-2+ subjects. A line of best fit was determined by simple linear regression (n=16).

We next interrogated the antibody response to the S2 subunit, where at a 1:50 dilution the SARS-CoV-2 positive subjects had an OD range of 1.3 to 3.1, while control subjects had an OD range of 0.1 to 1.07 (Figure 2B). Only one control subject had antibody binding that overlapped with the lower range of the SARS-CoV-2 subjects. We used an identical approach to evaluate the antibody response to RBD (Figure 2C). Here we saw a robust antibody response with a range in OD from 0.9 to 3.5 at a serum dilution of 1:50, and we observed no antibody binding above background in the control group. Multiple groups have observed a similar responses to the RBD subunit, indicating the specificity of the RBD antibody response ^33,34^, which could in part be due to the low level of conservation of the RBD amino acid sequences between SARS-CoV-2 and the HCoVs, which cause the common cold ^24,34^.

To further quantify differences in binding to the individual spike subunits we calculated the area under the curve (AUC) of the S1, S2, and RBD ELISA assays for each subject (Figure 2D-F, Table 3). Quantification of the AUC measures the antibody binding at multiple antibody concentrations quantifying a combination of avidity and specificity of the sera for each subject. When assessing binding to S1, the mean AUC of SARS-CoV-2 subjects was 2.61 +/− 1.38 and the mean AUC of controls was 0.71+/− 0.53 (p<0.0001; Figure 2D). Upon comparing the AUC for antibody binding to S2, we observed a mean AUC of 2.6 +/− 1.0 and 0.47 +/−0.22 (p<0.0001) SARS-CoV-2^+^ patients and controls, respectively (Figure 2E). Interestingly, when we assess binding to RBD, we observed a mean AUC of 3.8 +/−1.7 and 0.3 +/−0.07 (p<0.0001) showing minimal cross-reactivity from the negative subjects (Figure 2F). The RBD binding data matches recently described results ^33,34^, which show that the antibody response to RBD is specific to SARS-CoV-2 infection, with no known cross-reactivity from antibodies derived from endemic HCoV infection. Overall, we show that S1, S2, and RBD from spike are targeted by the human polyclonal response in all individuals from our cohort. Additionally, we observe potential cross-reactivity within the control group to the S1 domain outside of the RBD. This cross-reactivity is important to note for serological and vaccine evaluation, as using RBD as a target antigen may provide the most specific and sensitive test that results with fewer false-positives. Interestingly, this also highlights the potential for cross-reactive S1 antibodies to play a role in either protection or exacerbation of SARS-CoV-2 disease. It has been recently demonstrated that human mAbs generated after SARS-CoV infection were shown to cross-react and neutralize SARS-CoV-2 (^35^), while SARS-CoV infection generates a polyclonal antibody response that is able to bind spike from SARS-CoV-2 while not able to neutralize the virus ^36^. Furthermore, mechanisms aside from neutralization that are dependent on the Fc region of the antibody are capable of limiting viral infection ^15,37,38^.

As there was a broad dynamic range of antibody binding to spike subunits from our SARS-CoV-2^+^ subjects, we stratified the samples based upon days post qRT-PCR positive test, because days post symptoms was unavailable, to evaluate the potential role of time post infection on antibody binding and specificity. Samples were stratified into three groups: prior to day 18, day 21-26, and day 26-40 post qRT-PCR^+^ test. We compared the AUC values from the ELISA binding curves to S1, S2, and RBD over this time period and did not observe a role for time post infection on antibody binding and specificity with our limited sample set (Supplemental Figure 1 A-C). The heterogeneity of antibody binding has been observed in other patient cohorts ^33,34,39^.

Additionally, we quantified the relative binding of the polyclonal antibody response between different spike subunits to determine if the subunits were equally targeted by the antibody response. To this end, we evaluated the correlation of AUC between the S1, S2 and RBD subunits (Figure 2 G-I). When we compared the AUC of S1 and S2 we observed significant correlation (p=0.0055, r=0.6265) (Figure 2G). As expected, based on the location of RBD within S1, S2 AUC significantly correlated with RBD AUC (p=0.0100, r=0.5824), which may suggest epitopes for binding within S2 as well as RBD (Figure 2H). Based on the location of RBD within S1 we would anticipate correlation between their AUC values (Figure 2I), and indeed there is a significant correlation (p=0.0120, r=0.5676) that would suggest that either the majority of binding to S1 occurs within RBD, or that there are antibody epitopes throughout S1 that drive a robust antibody response. The ELISA antibody binding results indicate that all SARS-CoV-2^+^ patients within our cohort had antibodies which bound to each subdomain of the spike protein.

### Neutralization potential of sera from SARS-CoV-2^+^ cohort

Antibody neutralization is one mechanism of protection from severe viral disease. The mechanism of action of neutralizing antibodies often include the targeting of viral proteins that interact with the host receptor for entry or viral proteins required for fusion with host cell membranes (reviewed in^40^). For SARS-CoV-2, the multifunctional spike protein is required for entry and fusion. Specifically, the S1 domain contains RBD, which is responsible for binding the human ACE2 protein mediating entry ^41,42^, while S2 contains the fusion peptide ^43^. It has been shown by other groups that monoclonal antibodies targeting spike can block infection with SARS-CoV-2 and that natural infection of humans often produces neutralizing antibodies ^29–32,34,44,45^, which is thought to prevent subsequent COVID-19. However, the specificity of human polyclonal neutralizing antibodies against infectious SARS-CoV-2 is only now beginning to be understood.

To begin to understand the human polyclonal neutralizing antibody response we utilized a focus reduction neutralization tests (FRNT) (Figure 3A) based upon the assay we had developed for multiple emerging infectious diseases ^46–48^ and for SARS-CoV-2^49^ (Figure 3A). There are multiple advantages to the FRNT assay over pseudotype-virus assays and plaque assays, including the use of infectious virus that may better reflect heterogeneity in the conformational structure of the virion, quantitative measurement of the reduction of viral replication and spread as each foci diameter measured represents multiple cells, and finally the use of 96 well plates allowing for titers to be quantified using multiple technical and biological replicates. Overall, this assay allows for a rigorous and quantitative determination of antibody neutralization potential.

**Figure. 3.**
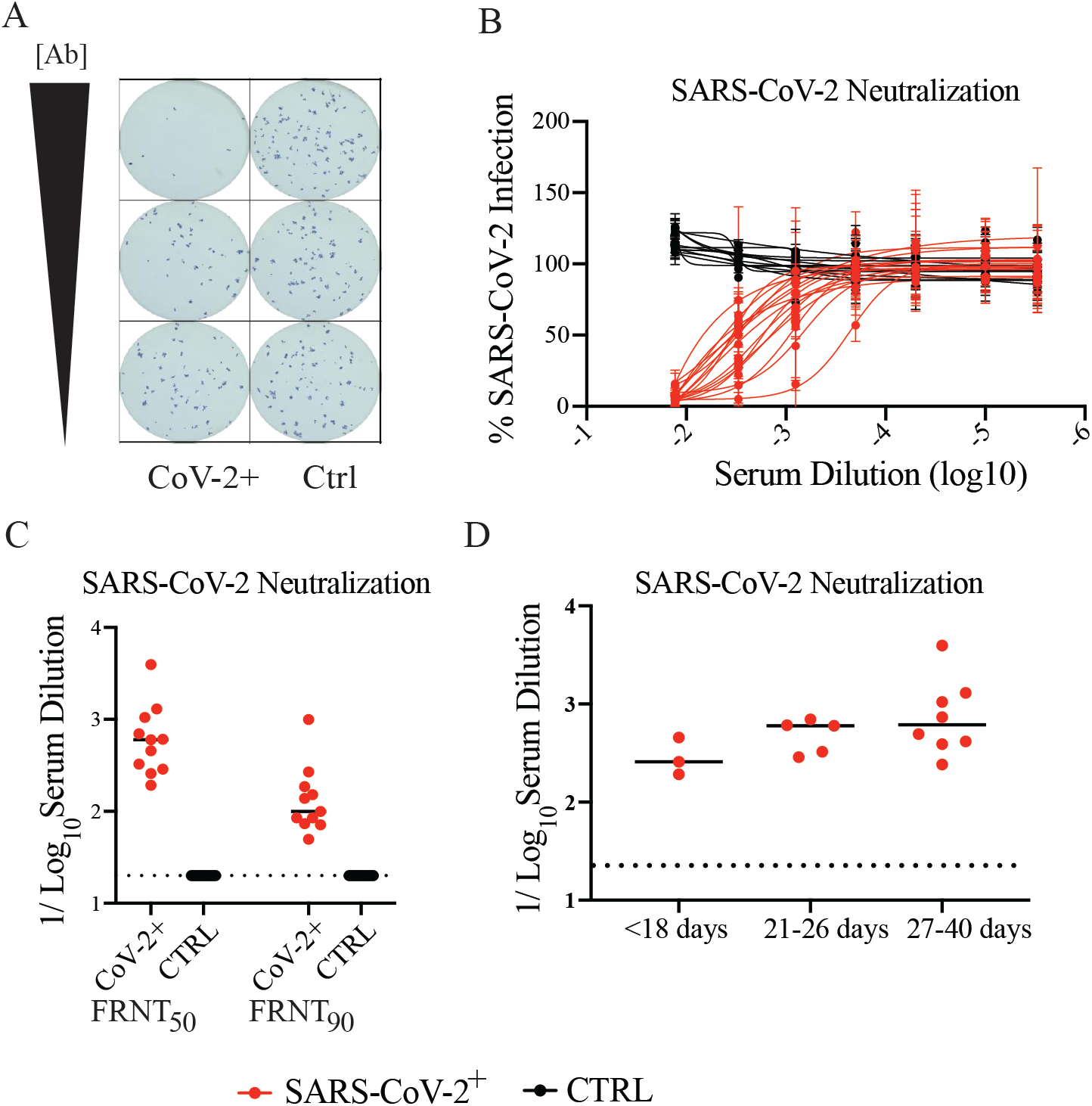
Neutralization potential of human sera against SARS-CoV-2 Neutralization of active SARS-CoV-2 by human sera was determined by Focus Neutralization Reduction Test (FRNT). (A) A representative section of a 96 well plate is shown for visualization of assay development. Increasing dilutions of neutralizing antibody are shown in column 1, while column is serial dilution of ctrl sera. Viral infected foci were stained with guinea pig-anti-SARS-CoV and developed. Foci equal to 2 cell diameter or larger are score as positive to quantify the neutralization of replicating virus. (B) Quantification of the FRNT assay is represented as a neutralization curve derived from non-linear regression analysis. Data are representative of 16 SARS-CoV-2+ subjects (red) and 14 controls (black). (C) FRNT_50_ and FRNT_90_ values were determined by non-linear regression analysis for SARS-CoV-2+ (n=16) and control (n=14) subjects. (D) FRNT_50_ values were stratified depending on time post positive qRT-PCR test. Groups were stratified into <18 days, 21-26 days, and 27-40 days post test.

Using the FRNT assay, we determined the concentration of patient sera required to neutralize SARS-CoV-2 infection. Based upon the antibody neutralization curve (Figure 3B), the serum dilution necessary to neutralize 50% of the virus (FRNT_50_) ranged from 1/53 to 1/4168 with a mean of 1/768 (Figure 3C). The serum dilution necessary to neutralize 90% of the virus (FRNT_90_) ranged from 1/50-1/995 with a mean of 1/200 (Figure 3C). Notably, the sera from SARS-CoV-2 patients in our cohort were capable of neutralizing infectious virus independent of day post positive test (Figure 3B-C, Table 2); while, sera from the majority of control subjects had no demonstrated antibody neutralization. One control subject, whose sera was cross-reactive in the S1/S2 ELISA binding assay demonstrated 10% SARS-CoV-2 neutralization potential at a 1/50 dilution, but further investigation of cross-neutralization is beyond the scope of this current study. Based upon the ability of the SARS-CoV-2 subjects to neutralize at least 90% of the virus, we show that the polyclonal antibody response has the breadth and specificity to completely neutralize SARS-CoV-2 infection. This would suggest that natural infection would be capable of controlling viral infection and limiting the potential of disease and transmission at the timepoints we assessed (Figure 3D). In animal model studies, hamsters have demonstrated that immune sera can protect from challenge^50^, although currently the mechanisms of that protection are unknown.

**Table 2.** FRNT_50_ and FRNT_90_ values determined by FRNT for SARS-CoV-2^+^ patients. The reciprocal serum dilution required to neutralize 50% (FRNT_50_) and 90% (FRNT_90_) of the virus was determined by non-linear regression analysis.

| **SAMPLE NO.** | **FRNT<sub>50</sub>** | **FRNT<sub>90</sub>** |
| --- | --- | --- |
| 1 | 1170.28 | 268.60 |
| 2 | 588.58 | 151.77 |
| 3 | 355.01 | 84.96 |
| 4 | 538.50 | 85.11 |
| 5 | 399.20 | 73.69 |
| 6 | 766.87 | 138.77 |
| 7 | 187.55 | 49.75 |
| 8 | 4338.40 | 995.01 |
| 9 | 299.76 | 99.60 |
| 10 | 264.48 | 71.43 |
| 11 | 1425.92 | 185.80 |
| 12 | 736.38 | 103.36 |
| 13 | 241.60 | 32.63 |
| 14 | 390.77 | 72.46 |
| 15 | 416.32 | 37.58 |
| 16 | 494.32 | 72.89 |

### Determination of the antigenic targets of human neutralizing polyclonal antibodies

To functionally determine which of the spike subunits are the main target of neutralizing antibodies, we performed a functional assay developed by the de Silva lab for use in flaviviruses ^51^. In this approach individual spike subdomains are linked to beads and are used to depleted sera in an antigen specific manor. In our studies his-tagged proteins are conjugated to cobalt coated magnetic beads and serum from SARS-CoV-2 subjects are incubated with the conjugated proteins. This allows a complex of antibody:antigen:bead to form and be pulled down by a magnet, leaving the serum depleted of that particular antibody specificity (Figure 4A). To understand the contribution of antibodies specific to each individual subunits, antibodies specific to each spike subunit, S1, S2, and RBD, were depleted from human polyclonal sera, and the antibody binding and neutralization potential of polyclonal sera after depletion was determined by ELISA and FRNT, respectively. Using the bead-based approach, sera from 10 patients were depleted for S1, S2, and RBD individually by sequentially incubating serum two times with protein coated beads.

**Figure. 4.**
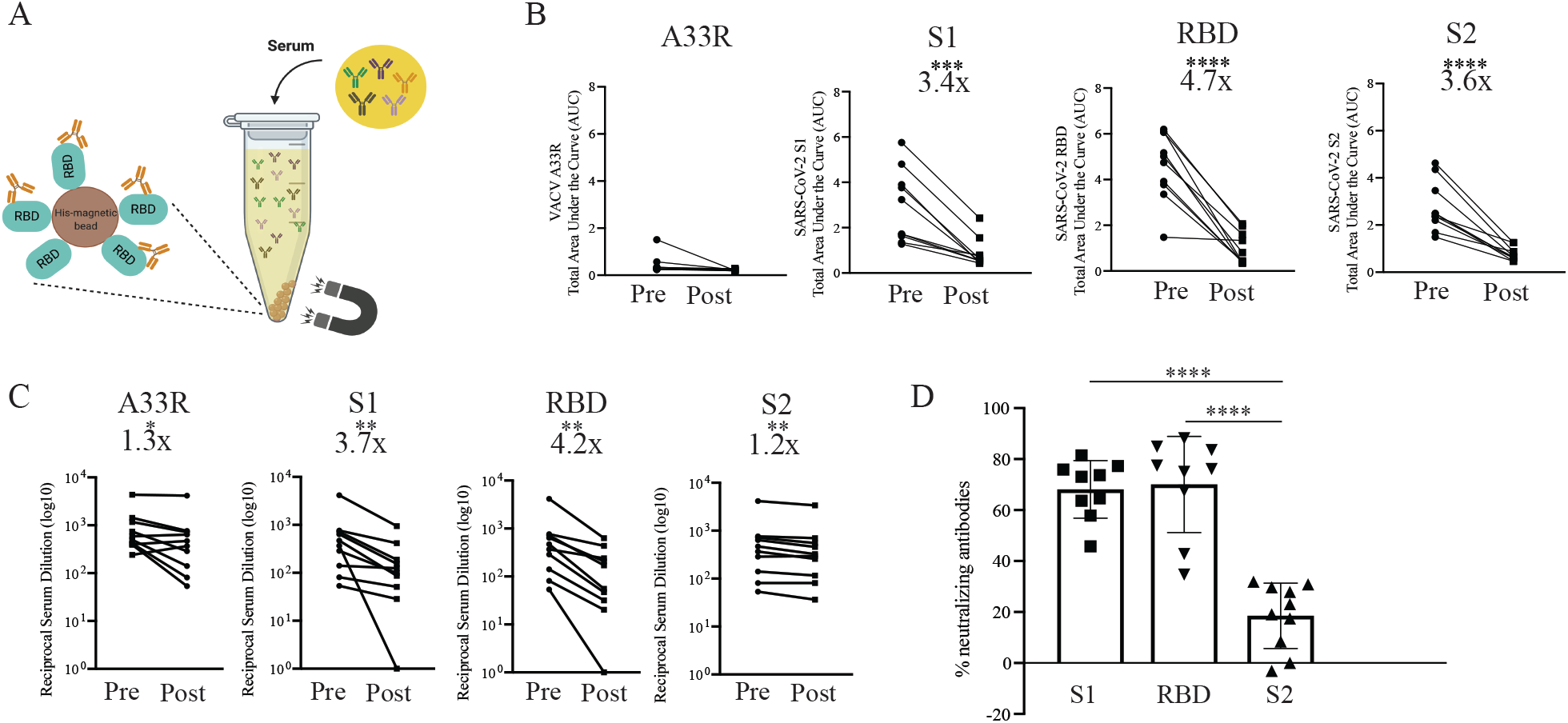
Antigen specific immune depletion of polyclonal sera and impact on SARS-CoV-2 neutralization potential. Immune depletions of antigen specific antibodies from human sera was accomplished in a his-specific magnetic bead-based approach, as depicted graphically (A). The efficiency of immune depletions for 10 SARS-CoV-2+ subjects was determined by ELISA and the area under the curve was calculated for (B) control protein (VACV A33R), S1, S2, and RBD. AUC values from pre and post depletion was paired for each subject to determine fold reduction in binding. (C) Reduction in FRNT50 post antibody depletion for VACV A33R, S1, RBD, and S2 was determined by FRNT. Similarly, FRNT50 values were paired pre and post depletion for each subject and statistical significance was determined by Wilcoxon matched-pairs rank test *(*p* = 0.0322), **(*p* = 0.0022), ***(*p* = 0.0002), and ****(*p* < 0.0001). (D) The percent of neutralizing antibodies that bind to each subunit of S was determined by the following equation: (1-(post depletion FRNT_50_/control depletion FRNT_50_))*100. Statistical significance was determined by Mann-Whitney test *(*p* = 0.0322), **( *p* = 0.0021), *** (*p* = 0.0002), ****( *p* <0.0001). Data are representative of n=10.

To quantify the effects of the antigen-specific antibody depletions, the AUC from ELISA binding curves pre and post depletion (Supplemental Figure 2, Table 3) were quantified, and the values were paired per subject (Figure 4B). After antigen specific depletions we observed significant reduction in spike subunit antibodies represented by a 3.4 (p=0.0005), 3.6 (p<0.0001) and 4.7 (p<0.0001) fold reduction in AUC binding to S1, S2, and RBD respectively (Figure 4B). Moreover, to confirm that depletion protocol did not impact SARS-CoV-2 neutralization we performed depletions with an irrelevant protein, VACV A33R (Figure 4B). The subunit depletion protocol significantly reduced the level of subunit specific antibody, which allowed us to evaluate the contribution of each individual subunit to the neutralizing antibody response.

**Table 3.**
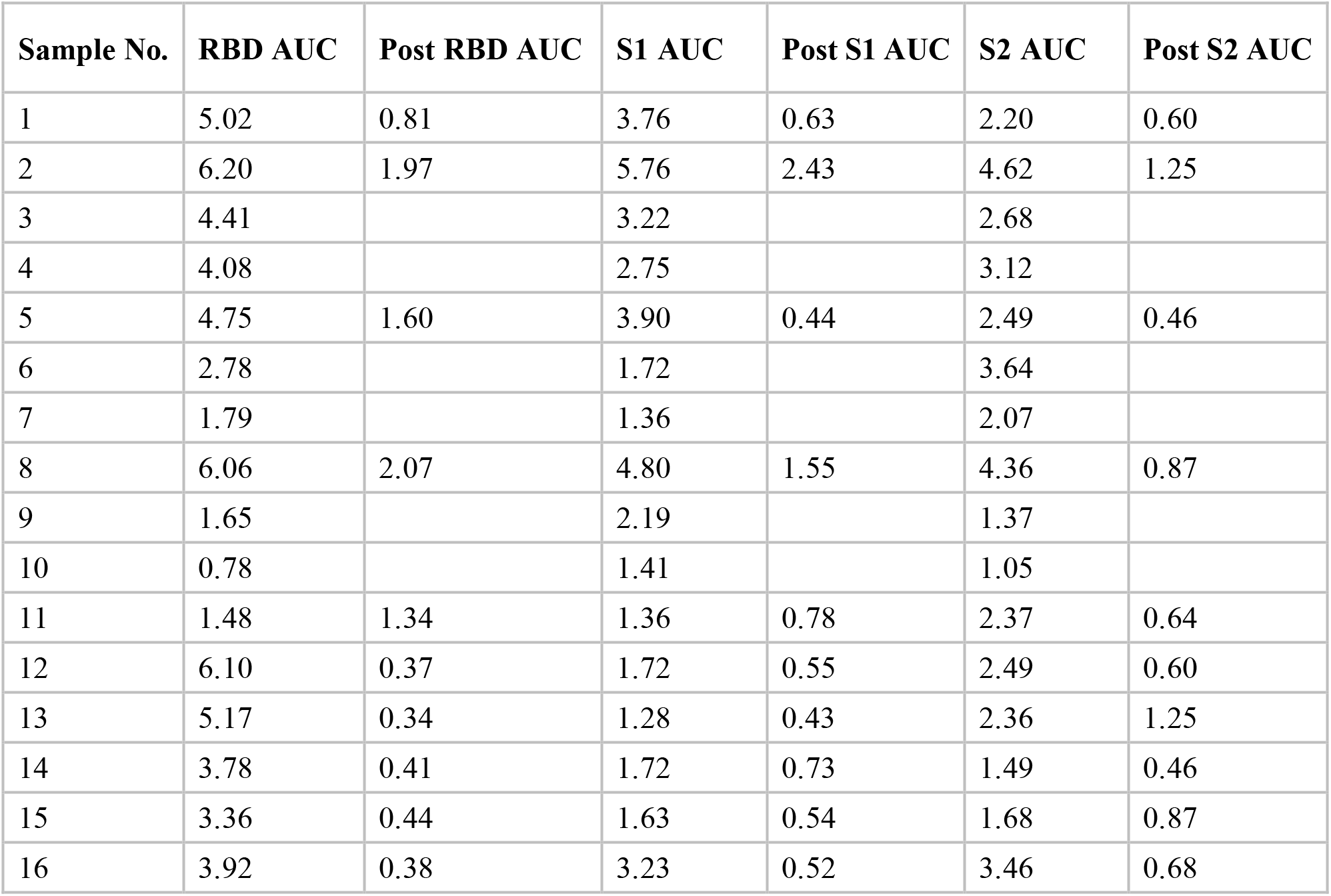
AUC values pre and post antigen specific depletion. Area under the curve (AUC) was calculated per subject pre and post depletion by ELISA assay.

| Sample No. | RBD AUC | Post RBD AUC | S1 AUC | Post S1 AUC | S2 AUC | Post S2 AUC |
| --- | --- | --- | --- | --- | --- | --- |
| 1 | 5.02 | 0.81 | 3.76 | 0.63 | 2.20 | 0.60 |
| 2 | 6.20 | 1.97 | 5.76 | 2.43 | 4.62 | 1.25 |
| 3 | 4.41 |  | 3.22 |  | 2.68 |  |
| 4 | 4.08 |  | 2.75 |  | 3.12 |  |
| 5 | 4.75 | 1.60 | 3.90 | 0.44 | 2.49 | 0.46 |
| 6 | 2.78 |  | 1.72 |  | 3.64 |  |
| 7 | 1.79 |  | 1.36 |  | 2.07 |  |
| 8 | 6.06 | 2.07 | 4.80 | 1.55 | 4.36 | 0.87 |
| 9 | 1.65 |  | 2.19 |  | 1.37 |  |
| 10 | 0.78 |  | 1.41 |  | 1.05 |  |
| 11 | 1.48 | 1.34 | 1.36 | 0.78 | 2.37 | 0.64 |
| 12 | 6.10 | 0.37 | 1.72 | 0.55 | 2.49 | 0.60 |
| 13 | 5.17 | 0.34 | 1.28 | 0.43 | 2.36 | 1.25 |
| 14 | 3.78 | 0.41 | 1.72 | 0.73 | 1.49 | 0.46 |
| 15 | 3.36 | 0.44 | 1.63 | 0.54 | 1.68 | 0.87 |
| 16 | 3.92 | 0.38 | 3.23 | 0.52 | 3.46 | 0.68 |

To measure the functional effect of S1, S2, and RBD antibody depletion on virus specific neutralization we evaluated post-depletion neutralization activity by FRNT (Supplemental figure 3). To confirm that the depletion protocol itself had no off-target effects on SARS-CoV-2 neutralization, a control depletion with VACV A33R was completed and neutralization pre and post depletion was measured. The control depletion had a minimal effect on the ability of the polyclonal sera to neutralize SARS-CoV-2 (Figure 4C: FRNT_50_:1.3 fold decrease; FRNT_90_:1.8 fold decrease). We then measured the antibody neutralization curves after depleting serum with S1, RBD or S2, and determined the serum dilution required to reduce infection by 50% (FRNT_50_) and 90% (FRNT_90_) (Table 4). To take into account the effects of the antibody depletion protocol, we compared the FRNT_50_ of the control depleted serum with the subunit depleted serum, and observed a 3.7, 4.2, and 1.2 fold reduction after S1, RBD, and S2 depletion, respectively (Figure 4C). Based upon the FRNT_50_ and FRNT_90_ values, the depletion of S1 and RBD significantly reduced virus neutralization (p=0.0020 and p=0.0020). This suggests that polyclonal antibody binding to the RBD domain of the spike protein represents the key target of neutralizing antibody to SARS-CoV-2 after natural infection. Since we observed a similar fold reduction after S1 and RBD depletion, it is likely that the majority of the neutralizing response is found within the RBD domain of S1. However, this is the average neutralizing antibody response, which is applicable to our cohort. When we evaluate changes in individuals, there are two patients that have a strong RBD neutralizing response, but also have a S2 specific neutralizing antibody response with 1.47, and 1.21 fold change after S2 depletion. Overall, these data demonstrate natural SARS-CoV-2 infection generates a robust anti-RBD polyclonal neutralizing antibody response with some individuals mounting a neutralizing antibody response to S2. We conclude that the polyclonal neutralizing antibody response to SARS-CoV-2 primarily targets receptor interactions (S1/RBD) in the majority of individuals.

**Table 4.** FRNT_50_ and FRNT_90_ values post antigen specific antibody depletions as determined by FRNT. The impact of subunit specific antibody depletion was determined by FRNT and the reciprocal serum dilution required to neutralize 50% (FRNT_50_) and 90% (FRNT_90_) of the virus was calculated by non-linear regression analysis.

| Sample No. | Post A33R FRNT<sub>50</sub> | Post A33r FRNT<sub>90</sub> | Post S1 FRNT<sub>50</sub> | Post S1 FRNT<sub>90</sub> | Post RBD FRNT<sub>50</sub> | Post RBD FRNT<sub>90</sub> | Post S2 FRNT<sub>50</sub> | Post S2 FRNT<sub>90</sub> |
| --- | --- | --- | --- | --- | --- | --- | --- | --- |
| 1 | 716.33 | 105.69 | 188.43 | 52.68 | 170.82 | 18.35 | 550.96 | 157.28 |
| 2 | 637.76 | 124.01 | 153.89 | 12.92 | 206.23 | 14.73 | 459.77 | 84.60 |
| 5 | 468.82 | 75.24 | 86.88 | 13.28 | 55.22 | 0.903 | 324.15 | 96.81 |
| 8 | 4168.40 | 485.91 | 946.97 | 30.74 | 627.75 | 4.89 | 3372.68 | 347.58 |
| 11 | 1009.49 | 97.75 | 413.74 | 15.13 | 436.49 | 19.23 | 698.32 | 131.18 |
| 12 | 286.70 | 42.54 | 98.62 | 2.04 | 47.01 | 2.99 | 295.68 | 40.60 |
| 13 | 365.90 | 32.34 | 1.00 | 13.65 | 238.95 | 0.64 | 257.14 | 23.47 |
| 14 | 53.39 | 21.50 | 28.61 | 17.38 | 1.00 | 15.22 | 36.38 | 14.71 |
| 15 | 80.78 | 60.53 | 51.07 | 9.02 | 20.15 | 17.00 | 80.78 | 29.05 |
| 16 | 140.51 | 48.05 | 124.72 | 19.67 | 31.72 | 10.04 | 115.98 | 26.83 |

To compare the relative neutralizing differences between spike domains, we normalized the data based upon FRNT_50_ values and represented the data as % subunit specific neutralizing antibodies. This allows us to calculate the percentage of neutralizing antibodies that bind to S1, S2, or RBD, while taking into account the impact of the depletion protocol, based on our control and subunit specific depletions (Figure 4D). Further confirming the paired FRNT data, 70% +/−19 and 68% +/−11% the highest percentage of neutralizing antibodies indeed bind to RBD and S1, suggesting a prevention in virus interaction with viral receptor maybe the dominant mechanism for antibody neutralization of SARS-CoV-2 after natural infection. Additionally, S2 has the lowest percentage 19% +/− 13% of S2 binding antibodies capable of neutralization, suggesting that viral fusion with host membranes is not a dominant target of the neutralizing antibody response to SARS-CoV-2 after natural infection, with our cohort of patients (Figure 4D). This data has been further represented as % binding neutralizing antibodies based on the pre depletion FRNT_50_ values (Supplemental Figure 3D). Overall, these data further confirm that a majority of neutralizing antibodies are targeted against the RBD within S1.

## Discussion

In this study we examined the antigenic targets of the SARS-CoV-2 IgG neutralizing antibody response that develop during natural infection. We quantified the immunodominance of anti-spike subdomain antibodies for binding by ELISA and neutralization activity by antigen specific depletion followed by a SARS-CoV-2 neutralization assay. To define the specificity of the antibody response during natural infection, we needed to understand the amino acid variation present in the currently circulating SARS-CoV-2 human isolates. Human SARS-CoV-2 isolates has a low frequency of amino acid variation within the spike protein, with the exception of the D614G mutation, allowing us to estimate that the majority of known isolates permit effective polyclonal antibody binding and neutralization. The human polyclonal antibody response recognizes three subdomains (S1, S2, and RBD) of the spike protein as evidenced by ELISA. Interestingly, we identified cross-reactive sera from SARS-CoV-2 naïve subjects to S1 suggesting conserved sequences in the S1 subunit of spike may impact non-neutralizing responses to SARS-CoV-2 as well as serological tests for SARS-CoV-2. Most importantly, our antigen-specific antibody depletion approach demonstrated that the RBD domain of the spike protein is responsible for 70% +/− 18.9% of the human polyclonal neutralizing antibody activity to spike after natural SARS-CoV-2 infection.

For the prioritization and selection of vaccine candidates for SARS-CoV-2, mechanisms of protection need to be defined. In order to understand the mechanisms of protection, we must dissect the specificity of the immune response to natural infection and vaccination. Several studies have begun to interrogate the binding characteristics of the polyclonal antibody response to SARS-CoV-2 and have identified the spike protein as a key target of the antibody response ^24,34,39^. Additional natural history studies have recently demonstrated that antibody binding to the RBD subdomain of the spike protein correlates with overall neutralizing antibody responses ^39,52^. Our study supports the conclusions that the SARS-CoV-2 spike protein is a key target of the human polyclonal antibody response after natural infection. Uniquely, our results functionally demonstrate that the RBD domain is the target of the majority of the neutralizing antibody response, while other studies have showed the correlation between RBD antibody binding and antibody neutralization. These data support the inclusion of the RBD domain in vaccine development and serological assay design.

The development of antibody based-therapeutics to combat recent emerging viruses including SARS-CoV, MERS-CoV, West Nile, Zika, and Ebola viruses relies on neutralizing activity as the correlate of protection. In order to determine which mAbs will be the most effective against SARS-CoV-2, sites present on the virion that are susceptible to antibody mediated neutralization need to be identified. Several groups have isolated neutralizing mAbs that target the N terminus of the S1 subdomain as well as the RBD of the spike protein, which have low nanomolar efficacy^14,29–31,44,53^. Our characterization of the polyclonal neutralizing antibody response indicates that all three subdomains of the spike protein have neutralizing antibody epitopes present, which can be targeted by the human antibody repertoire. The broad knowledge of this potential susceptibility would allow the targeted development of antibody cocktails that target all three spike subdomains, potentially preventing the development of antibody escape mutants that has already been documented *in vitro*.

Although our study shows that the dominant target of IgG neutralizing antibody response after natural SARS-CoV-2 infection is the RBD domain of the spike protein, we have evaluated a limited number (n=10) of patients by antigen-specific antibody depletion. There is the potential that immunodominance of the neutralizing antibody response may vary based upon a number of variables including viral load, co-morbidities including age and obesity, as well as genetic background. Additionally, we have only focused on the IgG response and it has been recently determined that the IgA antibody response can neutralize SARS-CoV-2 virus and the antigen specificity of that response could be different than the IgG response ^54^. Importantly, it has also been recently described that more than 90% of individuals who seroconvert generate detectible neutralizing antibody responses and that these IgG responses are indeed sustained for up to three months ^39,55^, which has the potential to protect against re-infection.

As our understanding of the immune response to natural infection improves it is important to begin to evaluate the correlates of protection beyond antibody neutralization, and investigate additional antibody mechanisms such as antibody dependent cellular cytotoxicity. As we detected antibodies targeted against S2 that are non-neutralizing these could provide a different mechanism of protection that may be valuable when considering vaccine design. There is also a strong T cell response established during natural infection ^56–59^, as well as a cross-reactive T cell response from potentially prior HCoV infection ^60,61^. Currently the role of the human T cell response to SARS-CoV-2 has only begun to be dissected.

Overall our study describes the polyclonal IgG response to SARS-CoV-2 from sera obtained from patients in a range of 14-40 days post positive qRT-PCR test. We focused on the relationship between antibody binding to the subdomains of spike and the neutralization capacity against infectious virus. We demonstrate that infection with SARS-CoV-2 results in an antibody response that results in a similar amount of IgG that targets spike subunits S1, S2, and RBD regardless of time post infection (Supplemental Figure 1). Furthermore, we show that this response results in a neutralizing antibody response by 14 days post positive qRT-PCR, as determined by FRNT (Figure 3B). Finally, using a bead-based immune depletion approach, we show that the highest percentage of neutralizing antibodies against SARS-CoV-2 bind to the receptor binding domain (RBD) (Figure 4D) that directly interacts with human ACE2. These findings are important in the further development and prioritization of therapeutics and vaccine development.

## Materials and Methods

### Ethics statement

The serum and plasma samples used for this study were collected at Case Western Reserve University (IRB:02-17-03), University of Puerto Rico (IRB: Pro00043338) and Saint Louis University (IRB: 26646, 27790). All patients were diagnosed with acute SARS-CoV-2 infection by qRT-PCR. All collection, processing and archiving of human specimens was performed under approval from the University Institutional Review Board.

### Viruses and cells

SARS-CoV-2 was obtained from Biodefense and Emerging Infections (BEI) Research Resources Repository and passaged once in Vero E6 cells purchased from American Type Culture Collection (ATCC CCL-81).

### Recombinant Proteins

Vaccinia Virus A33R Protein with C-Terminal Histidine Tag, was obtained through BEI Resources, NIAID, NIH: NR-2623. SARS-CoV-2 (2019-nCoV) Spike S1-His Recombinant Protein (40591-V08B1) and SARS-CoV-2 (2019-nCoV) Spike S2 ECD-His Recombinant Protein (40590-V08B) were obtained from Sino Biological. SARS-CoV-2 Spike protein (RBD, His Tag) (Z03483) was obtained though GenScript. Validation of proper protein folding was completed by mAb ELISA and size was done by SDS-PAGE.

### SARS-CoV-2 Sequence Analysis

Full length SARS-CoV-2 spike sequences derived from human isolates were identified and downloaded from GISAID. Sequences were trimmed and aligned with Geneious 7 and single amino acid variants were identified and quantified using the analyze sequence variation tool of the NIAID Virus Pathogen Database and Analysis Resource (VIPR) and shannon entropy calculations. The resulting amino acid changes were mapped onto the cryoEM structure of SARS-CoV-2 spike (6VSB) using the consurf.py script. Display of the aa variation on the cryo-em structure was completed using pymol v2.40.

### Enzyme Linked Immunosorbant Assay (ELISA)

Binding of human polyclonal sera to recombinant SARS-CoV2 proteins was determined by ELISA. Briefly, MaxiSorp (ThermoFisher) plates were coated with 50uL of a 1ug/mL mixture of recombinant protein in carbonate buffer (0.1M Na2CO3 0.1M NaHCO3 pH 9.3) overnight at 4°C. The next day plates were blocked with blocking buffer (PBS + 5%BSA + 0.5% Tween) for 2 hours at room temperature and washed 4x with wash buffer prior to plating of serially diluted polyclonal sera. Sera was incubated for 1 hour at room temperature in the ELISA plate, washed 4x with wash buffer, followed by addition of goat-anti-human IgG HRP (Sigma) conjugated secondary (1:5000) for 1 hour at room temperature. The plate was washed again 4x with wash buffer and the ELISA was visualized with 100uL of TMB enhanced substrate (Neogen Diagnostics) and placed in a dark space for 15 minutes. The reaction was quenched with 1N HCl and the plate was read for an optical density of 450 nanometers on a BioTek Epoch plate reader. Total peak area under the curve (AUC) was calculated using GrapPad Prism 8.

### Antigen Specific Antibody Depletions

Antigen specific antibodies were depleted in a bead-based approach using Ni-NTA Magnetic beads (Thermo Scientific) as described (^62^). SARS-CoV2 his tagged proteins or VACV his tagged protein (control depletion) were conjugated to the his-specific magnetic beads as suggested by manufacturer’s protocol. Briefly, 1mg of beads were washed with equilibration buffer followed by addition of 50ug of protein diluted in equilibration buffer. After addition of protein, the tube was rotated end over end for 1 hour at 4°C. The beads were collected on a magnetic stand and washed twice with wash buffer followed by separation into two tubes of 200μL each. Next, the human sera were diluted in tissue culture sterile PBS and placed into the first tube of beads and incubated end-over end at 37°C for 1 hour. Once again, the beads were collected with a magnetic stand, the supernatant was removed and transferred into the second tube for another end-over-end incubation at 37°C for 1 hour. After incubation the beads were collected, and the supernatant was removed and placed at 4°C for subsequent ELISAs and FRNTs. After ELISAs and FRNTs, the percentage of CoV-2-binding antibodies was calculated by dividing the depleted IgG titers by the control-depleted IgG titers. The percentage of neutralizing antibodies was calculated by % subunit binding neutralizing antibodies = ((1−(FRNT50 post depletion)/(FRNT50 post control depletion))*100.

### Focus Reduction Neutralization Test

Four-fold serial dilutions of human serum were mixed with ~100 focus-forming units (FFU) of virus, incubated at 37°C for 1 h, and added to Vero WHO monolayers in 96-well plates for 1 h at 37°C to allow virus adsorption. Cells were overlaid with 2% methylcellulose mixed with DMEM containing 5% FBS and incubated for 24 hours at 37°C. Media was removed and the monolayers were fixed with 5% paraformaldehyde in PBS for 15 min at room temperature, rinsed, and permeabilized in Perm Wash (PBS, 0.05% Triton-X). Infected cell foci were stained by incubating cells with polyclonal anti-SARS Guinea Pig sera for 1 h at 37°C and then washed three times with Perm Wash. Foci were detected after the cells were incubated with a 1:5000 dilution of horseradish peroxidase-conjugated goat anti-guinea pig IgG (Sigma) for 1 h. After three washes with Perm Wash, staining was visualized by addition of TrueBlue detection reagent (KPL). Infected foci were then enumerated by CTL Elispot. FRNT curves were generated by log-transformation of the x axis followed by non-linear curve fit regression analysis using Graphpad Prism 8.

## Supporting information

Spike_msa

VIPR and CONSURF tables

## Acknowledgments

We thank the volunteers for providing there time and samples for these studies.

## Funding

This work was supported by Saint Louis University COVID-19 research Seed Funding to awarded to AKP and awarded to JDB, and supported by National institutes of Health grant F31 AI152460-01 from the National Institute of Allergy and Infectious Diseases (NIAID) awarded to MH. The Puerto Rico Science, Technology and Research Trust supported research reported in this work under agreement number 2020-00272 to AME and CAS.

## Author contributions

TLS, ETS, MH, BTG, AME, PP, CC, CAS, AKP, and JDB conceptualized the work and TLS, ETS, MH, DFH, SLG, AKP, and JDB wrote and edited the manuscript. Design and execution of the FRNT assays was completed by ETS, TLS and JDB. Design and execution of the ELISA assays and sera depletion studies were completed by TLS. All authors reviewed and approved the final version of the manuscript.

## Competing Interests

The authors declare no competing interest

## Data and materials availability

All data is present on BioRX.

## Supplementary Materials

**Supplemental Figure 1**. Antibody binding and specificity to spike sub-domains over time.

To evaluate a potential role for time post infection on antibody binding and specificity, subjects were stratified into three groups based on time of sample collection post positive SARS-CoV-2 test (14-16, 21-23, and 26-40 days post-test (d.p.t)). The AUC values derived from the binding curves were stratified based on the criteria and differences in binding were assessed for (A) S1, (B) S2, and (C) RBD. Data is representative of n=3 for 14-16 d.p.t, n=4 for 21-23 d.p.t, and n=9 for 26-40 d.p.t.

**Supplemental Figure 2**. ELISA binding curves post depletion.

Binding of polyclonal IgG from SARS-CoV-2 qRT-PCR positive subjects to recombinant SARS-CoV-2 proteins pre and post antigen specific antibody depletion was determined by indirect ELISA. Binding curves derived from optical density at A450 for each subject are shown for (A) S1, (B) S2, (C) RBD, and (D) VACV A33R to determine the quantity of decreased binding. Red lines represent SARS-CoV-2^+^ subjects pre depletion and the black lines represent SARS-CoV-2 + subjects post depletion. Data are representative of n=10 for pre and post depletion.

**Supplemental Figure 3**. FRNT curves post depletion.

Neutralization of active SARS-CoV-2 by polyclonal sera from SARS-CoV-2^+^ subjects pre and post antigen specific antibody depletion was determined by Focus Neutralization Reduction Test (FRNT). Neutralization curves are depicted for (A) post S1 depletion, (B) post S2 depletion, (C) post RBD depletion, and (D) post VACV A33R depletion. Red lines represent SARS-CoV-2^+^ subjects pre depletion and the black lines represent SARS-CoV-2 + subjects post depletion. Data are representative of n=10 for pre and post depletion. (E) The percent of neutralizing antibodies that bind to each subunit of S was determined by the following equation: (1-(post depletion FRNT_50_/pre depletion FRNT_50_))*100. Statistical significance was determined by Mann-Whitney test *(*p* = 0.0322), **(*p* = 0.0021), *** (*p* = 0.0002), ****(*p* <0.0001). Data are representative of n=10.

### Supplemental Data files S1-S2

Supplemental file 1: 4065 muscle alignment

Supplemental file 2: VIPR and CONSURF output tables

## Notes

### Competing Interest Statement

The authors have declared no competing interest.

