## Supplementary figures and images for "The receptor binding domain of SARS-CoV-2 spike is the key target of neutralizing antibody in human polyclonal sera"

### Supplemental Figure 1

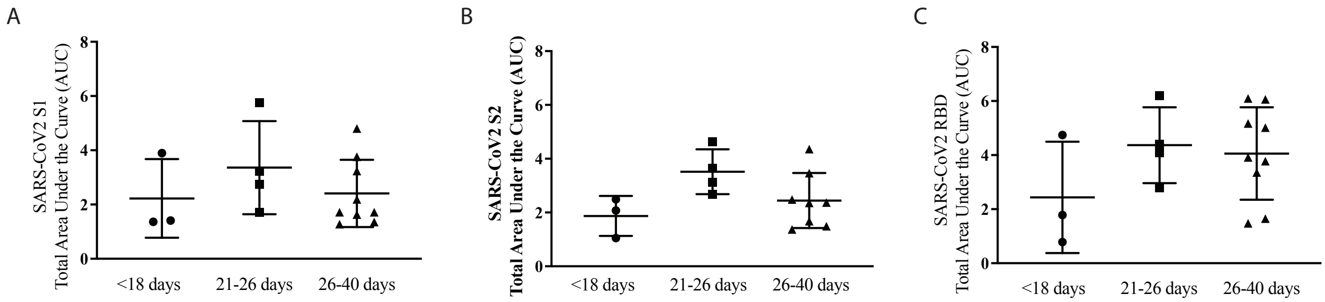


Supplemental Figure 1

### Supplemental Figure 2

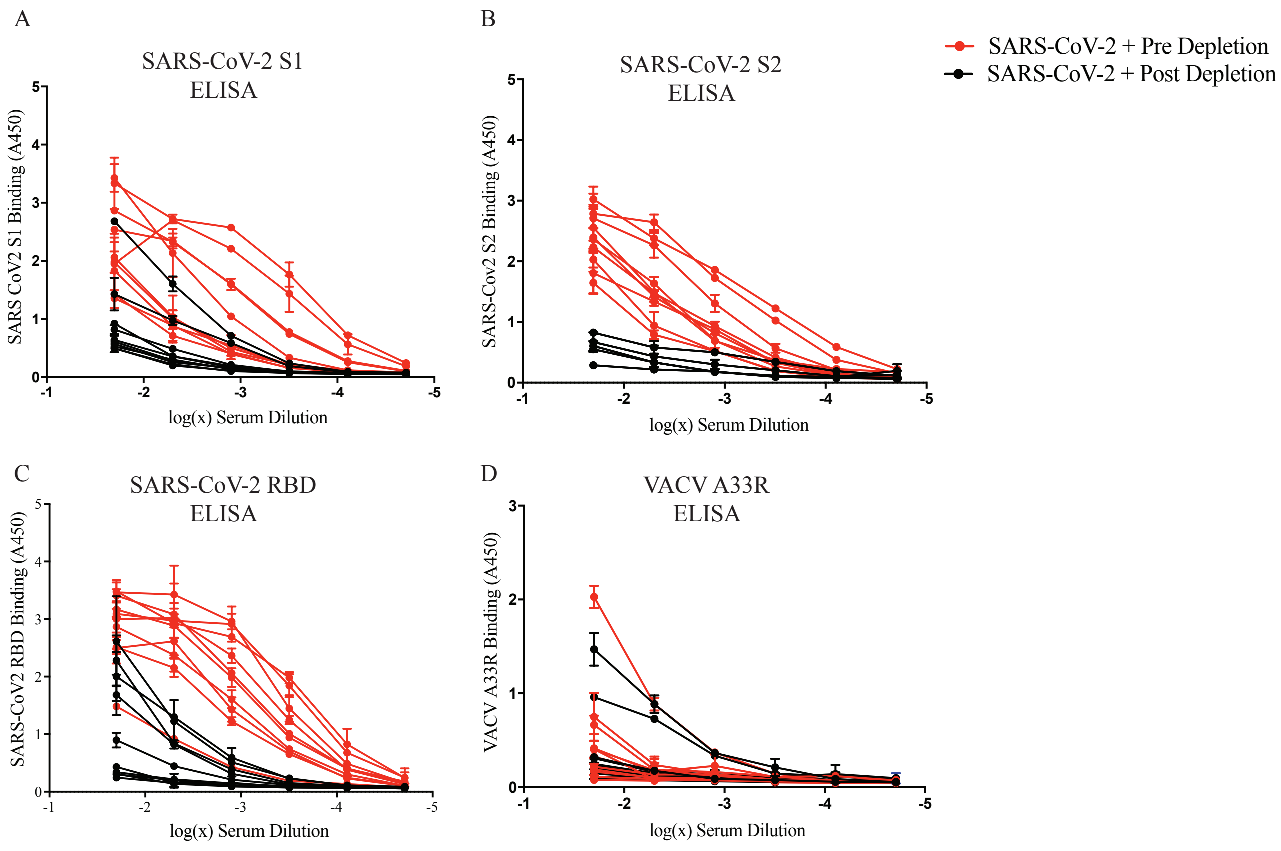


Supplemental Figure 2

### Supplemental Figure 3

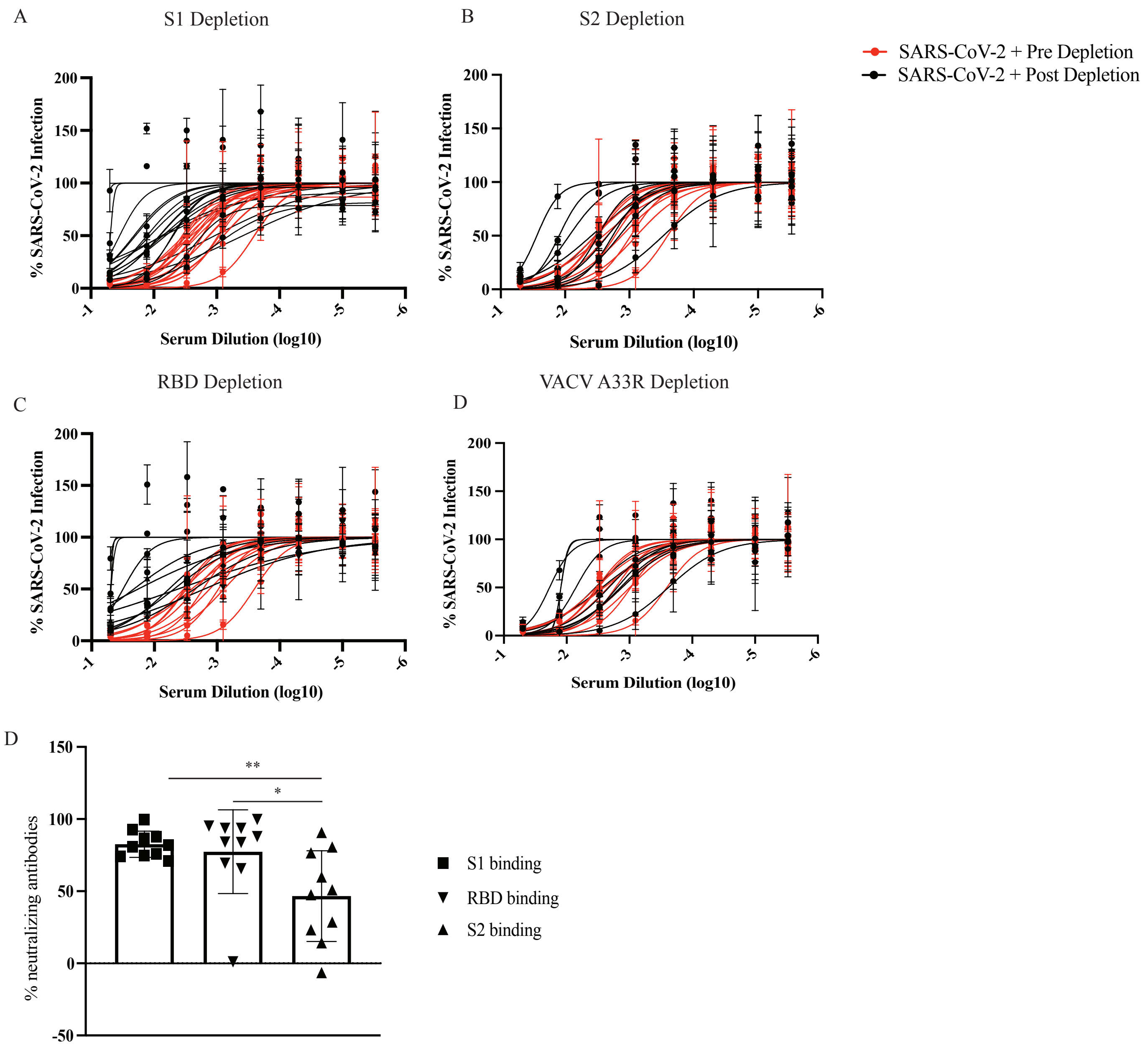


Supplemental Figure 3
